# Enhanced cytokine responsiveness in natural killer cells from a pilot cohort of uninfected seronegative women exposed to hepatitis C virus contaminated anti-D immunoglobulin

**DOI:** 10.1101/747436

**Authors:** M. W. Robinson, C. Keane, M. Needham, G. Roche, E. Wallace, J. Connell, C. F. de Gascun, A. Naik, L. J. Fanning, C. Gardiner, D. D. Houlihan, C. O’Farrelly

## Abstract

**Background:** Some people exposed to hepatitis C virus (HCV) appear to be capable of preventing infection in the absence of detectable antibody responses. These ‘exposed seronegative (ESN)’ people appear naturally resistant to HCV infection. Here, we aimed to examine innate immune mechanisms in ESN individuals amongst rhesus negative Irish women exposed to HCV via contaminated anti-D immunoglobulin between 1977-79 and 1991-94.

**Methods:** A total of 16 ESN individuals were recruited, along with 9 age- and gender-matched healthy controls. All tested negative for HCV-specific antibodies using conventional diagnostic assays. Peripheral blood cells were analysed for presence of adaptive immune response markers, innate immune responsiveness and natural killer cell phenotype and function.

**Results:** The innate immune cell profile of ESN women in the present study was characterised by a significant decrease in monocyte frequency and elevated levels of interleukin-8 and -18 compared to age- and gender-matched healthy controls. NK cells from ESN women had normal expression of NK cell receptors but increased IFNγ-production upon cytokine and target cell stimulation as well as enhanced natural killer (NK) cell STAT3 phosphorylation in response to Type I IFN.

**Conclusions:** We describe for the first time ESN individuals amongst Irish women with past exposure to HCV via contaminated anti-D immunoglobulin. NK cells from these ESN individuals are more responsive to cytokine signalling compared with age- and gender-matched controls. Human ESN cohorts can provide unique insights into the biological mechanisms associated with antigen-independent natural resistance to viral infection.

## Introduction

Hepatitis C virus (HCV) infection is a leading cause of liver disease and liver failure, often requiring transplantation. The global prevalence of HCV viremia is estimated to be 70 million(1), and it is spread via blood-to-blood contact, predominantly within persons who inject drugs (PWID). However, HCV is also spread within medical settings in instances where there is a breakdown of infection prevention procedures. Following exposure, approximately three quarters of acutely infected individuals will go on to develop chronic infection(2). The remaining acutely infected individuals are able to clear HCV within 6 months, and have detectable HCV-specific antibody responses but no detectable HCV RNA; these individuals are referred to as spontaneous resolvers.

In addition to the aforementioned outcomes, some individuals exposed to HCV lack diagnostic signs of acute infection. These individuals, referred to as exposed seronegative (ESN), appear to evade infection without generating antibody responses detectable using conventional anti-HCV diagnostic assays. These individuals are identified by their history of HCV exposure, and the absence of diagnostic evidence of infection. ESN individuals were first characterised as uninfected individuals with HCV-infected family members(3, 4) or uninfected individuals with occupational exposure to HCV(5–7). ESN individuals have also been identified in several PWID populations. Study subjects shared needles with known HCV infected individuals, yet despite this risk factor, failed to become infected or develop HCV-specific antibody responses detectable by standard diagnostic assays(8–10).

It is likely that the ESN phenotype arises due to different biological and immunological mechanisms. These could include defective viral entry, effective intrinsic or innate immunity (11–15), HCV-specific T cell responses(3–10), and HCV strain-specific neutralising antibodies, which are not detected by standard diagnostic assays(16). These mechanisms may act individually or in combination to prevent the establishment of HCV infection (Figure 1).

**Figure 1:**
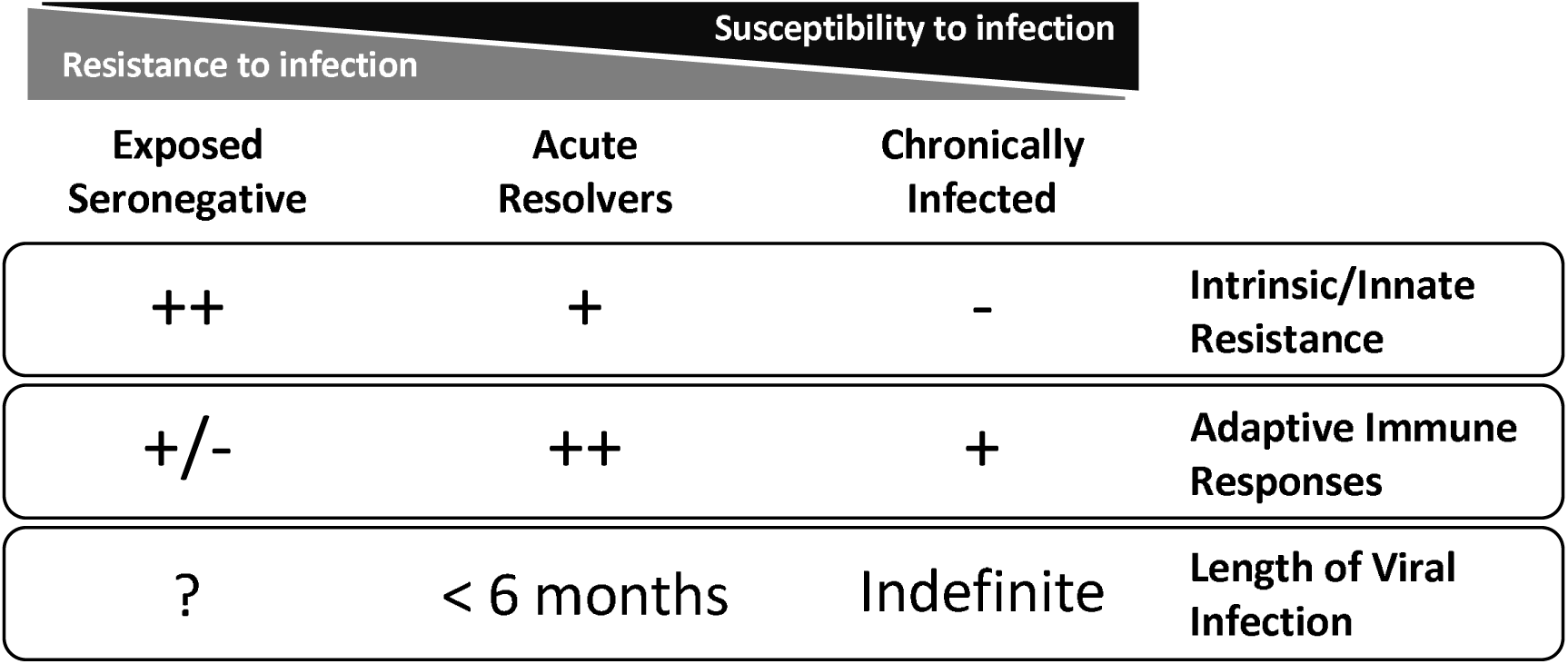
The spectrum of outcomes following hepatitis C virus exposure vary depending on a host’s resistance to virus infection. Exposed seronegative individuals may possess innate/intrinsic resistance to HCV infection and a proportion of ESN individuals possess adaptive HCV-specific immune responses not detected by current diagnostic assays. Acute resolution occurs within 6 months of exposure and is associated with rapid and robust adaptive HCV-specific immune responses. Adaptive immune responses are present in chronically infected individuals, although these are unable to clear the viral infection.

It is known that ESN individuals are more likely to carry genetic polymorphisms in the host viral entry co-receptor claudin-1(17), to lack the protective *IFNL3* genotype that is associated with spontaneous resolution and treatment sustained virological response (SVR)(18), and are more likely to be homozygous for a *KIR2DL3/HLA-C* allotype, associated with enhanced natural killer (NK) cell suppression of HCV replication(19). However, genetic analyses are unlikely to reveal functional mechanisms of resistance that involve several biological processes. Observational studies of ESN individuals, spontaneous resolvers, and chronically infected individuals, have identified hyper-responsive dendritic cell populations in ESN individuals(11) as well as enhanced NK cell function(6,12–15).

In Ireland, two single-source HCV outbreaks, the first between 1977 and 1979(20) (genotype 1b HCV), and the second between 1991 and 1994(21) (genotype 3a HCV), provide an opportunity to study ESN individuals without the co-morbidities seen in PWID. The 1977-1979 outbreak was traced back to a single donor, from whom 12 batches of potentially infectious anti-D immunoglobulin were prepared, and for which the infectivity rates have been published(22). The 1991-1994 outbreak was traced back to a different single donor. This outbreak was associated with significantly lower rates of infectivity, likely due to the relatively low level of HCV RNA present in the contaminated anti-D immunoglobulin preparations.

These Irish HCV outbreaks provide a unique research opportunity for three reasons: 1) the date of exposure to potentially infectious anti-D immunoglobulin for all recipients is known, and hence time since exposure can be calculated; 2) all recipients were potentially exposed to the same HCV strain (and the source strain from the original donor is known); and 3) all recipients were female of a similar age and health status, with few co-morbidities. We aimed to study innate immune responses in recipients of contaminated anti-D immunoglobulin who tested negative for anti-HCV antibodies upon screening and had no evidence of HCV infection. Understanding the biological mechanisms that contribute to resistance against HCV infection could provide novel avenues towards therapeutically manipulating host resistance to viral infection and future vaccination strategies.

## Methods

### Study recruitment

Seronegative individuals who received a potentially contaminated anti-D immunoglobulin product during either of the two Irish HCV outbreaks (HCV genotype 1b in 1977-1979 and HCV genotype 3 in 1991-1994) were invited to participate in the study. These seronegative individuals had self-reported to an Irish HCV patient support group as uninfected recipients of contaminated anti-D immunoglobulin. Age- and gender-matched control individuals who did not receive contaminated anti-D immunoglobulin were recruited at Trinity College Dublin, Ireland (n=9). These individuals were not matched for number of pregnancies or Rhesus blood group. As an additional control for the clinical testing and for T cell ELISpot experiments, a cohort of seropositive recipients of contaminated anti-D immunoglobulin was recruited through St. Vincent’s University Hospital, Ireland. This seropositive group (n=10) included 5 spontaneous resolvers and 5 individuals who achieved sustained virological response (SVR) following therapy. All seropositive individuals were negative for circulating viral RNA, in order to eliminate the direct effect of the active virus replication on immune responses(23). Research participants provided informed written consent in line with the Declaration of Helsinki, provided a clinical history, and blood samples for viral testing, biochemistry, immunology, full blood counts, and peripheral blood mononuclear cells (PBMC) extraction. Study protocols were approved by the St. Vincent’s University Hospital Research Ethics Committee and the Trinity College Dublin Faculty of Health Sciences Research Ethics Committee.

### Screening for adaptive immune markers of viral infection

IFNγ ELISpots were performed using the MABTECH Human IFNγ ELISpot kit (MABTECH AB, Nacka Strand, Sweden; 3420-2A), as per manufacturer’s recommendations. PBMCs were stimulated using: the ProMix™ HCV Peptide Pool (ProImmune Limited, Oxford, UK; PX-HCV-E); three over-lapping 13 to 19-mer peptide pools (obtained from BEI Resources, NIAID, NIH; Supplementary Table 1) spanning NS3, NS4A, and NS4B from either HCV genotype 1b (J4; GenPept: AAC15722) or genotype 3a (K3a/650; GenPept: BAA06044), as appropriate for the exposure; the ProMix™ CEF Peptide Pool containing immunodominant epitopes from influenza, CMV and EBV (ProImmune Limited; PX-CEF-E); 5μg mL^-1^ concanavalin A (Sigma-Aldrich; c5275); and DMSO as a negative control. All peptides were dissolved in sterile DMSO (Sigma-Aldrich, Arklow, Ireland; D2650) and used at a final concentration of 3μg mL^-1^. Plates were read and analysed using an AID EliSpot Reader (AID GmbH, Straßberg, Germany).

Serum samples were screened for HCV sero-reactivity using the ARCHITECT Anti-HCV assay (Abbott, Wiesbaden, Germany), Epstein Barr virus (EBV) sero-reactivity using the LIAISON® VCA IgG assay (DiaSorin S.p.A., Saluggia, Italy) and cytomegalovirus (CMV) sero-reactivity using the ARCHITECT CMV IgG assay (Abbott), as per manufacturers’ recommendations. To screen for antibodies targeting HCV glycoproteins, HCVpp-H77 neutralisation assays were performed as described previously(24), using the phCMV-ΔC/E1/E2 H77 clone kindly provided by François-Loïc Cosset, INSERM, France. Plasma was combined into three pools, one containing 9 samples from age- and gender-matched healthy controls, and two ESN pools each containing 8 samples, and IgG was purified using the Ab SpinTrap and Buffer Kit (GE Healthcare Life Sciences, Little Chalfont, UK). HCVpp-H77 were mixed with IgG preparations at concentrations from 0.014 mg mL^-1^ to 0.400 mg mL^-1^ and incubated for 1 hour at 37°C. This mixture was added to the Huh7 cells and after 72 hours the media was removed from the cells and 50µL of Cell Culture Lysis Reagent (Promega, Madison, WI, USA) was added for 15 minutes. The lysate was mixed with 50 µL of Bright-Glo substrate (Promega, Madison, WI, USA) and luciferase activity was measured in relative light units (RLUs) in a GloMax system (Promega). The percentage infectivity was calculated as 100% x (1-HCVppRLUtest/HCVppRLUcontrol). Each sample was tested in triplicate in two independent experiments.

### Analysis of immune cell frequency and IFN*α* responsiveness

Phosphorylated (p)STAT1 and pSTAT3 analysis was carried out using a stain-fix-permeabilise-stain protocol. Whole blood was stained for surface markers using antibodies detailed in Supplementary Table 2 and were then left untreated or stimulated with 1000 IU/mL IFN-α (Roferon-A, Roche Pharmaceuticals) for 30 minutes. After the stimulation period, cells were fixed immediately by adding Lyse/Fix Buffer (BD Bioscience) and then permeabilized for 30 min on ice using Perm Buffer III (BD Bioscience). Intracellular staining for pSTAT1 (pY701) and pSTAT3 (pY705) was carried out at RT for 60 min, then cells were washed and acquired immediately using a Canto II (BD Bioscience). Data were analysed using FlowJo software.

### Analysis of NK cell phenotype and function

Multicolour flow cytometry was used to analyse NK cell expression of the maturation marker CD57 and the NK cell receptors NKG2A, NKG2D, NKp30, NKp44 and NKp46. phenotype. PBMCs were stained with fluorescently labelled monoclonal antibodies as detailed in Supplementary Table 2. Dead cell exclusion was carried out using Near IR LIVE/DEAD® Fixable Dead Cell Stains (Thermo Fisher Scientific), and staining was performed with FcR Block (BD Biosciences) and BD Horizon™ Brilliant Stain Buffer (BD Biosciences). Data were acquired on a LSR Fortessa (BD Biosciences) and analysed using FlowJo V10.0.8 (FlowJo, LLC., Ashland, OR, USA).

IFNγ production and cell-surface CD107a expression, a marker of NK cell degranulation, in response to cytokine stimulation and co-incubation with 721.221 HLA-negative target cells were measured. PBMCs were seeded in round bottom 96-well plates and stimulated at 37°C for 16 hours with interleukin (IL)12 at 30ng mL^-1^, IL2 at 500 Units mL^-1^, IL15 at 100ng mL^-1^ (Miltenyi Biotec), and IL18 at 25ng mL^-1^ (R&D Systems Minneapolis, CA, USA). PBMCs were co-incubated with 721.221 cells and anti-human CD107a for 4 hours, with the addition of Golgistop and Golgiplug (BD Biosciences) after 1 hour. Samples were surface stained and intracellular staining was performed using the FoxP3 staining buffer set (eBioscience, San Diego, CA, USA) using anti-human IFNγ (as detailed in Supplementary Table 2).

### Cytokine Array

Circulating inflammatory cytokines from plasma samples were measured using the BioLegend LEGENDplex™ Human Inflammation Panel (13-plex) as per the manufacturer’s instructions. The stained LEGENDplex™ beads were acquired on a FACS Canto (BD Biosciences) and data were analysed using the LEGENDplex™ Data Analysis Software version 7.0 (BioLegend). Heat maps were generated using R version 3.2.1.

### Statistical Analysis

All statistical analyses were performed using GraphPad Prism version 5.01 for Windows (GraphPad Software, San Diego, CA, USA). For two group comparisons, two-tailed unpaired t-test or Mann-Whitney test were used. For categorical variables Chi-Square Test was used. For all analyses *P*<0.05 was considered statistically significant.

## Results

### Cohort Recruitment

Sixteen self-reported ESN women were recruited into this pilot study (Table 1). The batch number of anti-D immunoglobulin received could be identified for 15 women. Published infectivity rates for the 1977-1979 outbreak indicate that 6 of the 12 known contaminated batches accounted for more than 98% of all chronic infections (batch numbers 237, 238, 245, 246, 250, and 252). The rate of recipient sero-positivity amongst these 6 batches ranged from 30-78% (compared with 0-11% for the other 6 batches), and the rate of chronic infection ranged from 10-43% (compared with 0-3% for the other 6 batches)(22). In the present pilot study, a total of 3 ESN individuals received anti-D immunoglobulin from a batch with a high infectivity rate, while 12 ESN individuals received anti-D immunoglobulin from a batch with a low infectivity rate (either from the 1977-79 or the 1991-94 outbreaks). Due to the small size of the current study, individuals in the ESN group were not segregated by batch.

**Table 1.** Cohort demographics. Data for age are presented as mean with the 95% confidence interval in brackets. *Amongst the ESN individuals, one individual was exposed during both the 1977-79 and 1991-94 outbreaks.

|  | Matched Healthy Controls (HC) | Exposed Seronegative (ESN)* |
| --- | --- | --- |
| **n** | 9 | 16 |
| **Age (Years)** | 56.0 (52.0 – 60.0) | 57.8 (52.4 - 61.3) |
| **No. Exposed 1977-79**<br>**(HCV genotype 1A)** | N/A | 5 |
| **No. Exposed 1991-94**<br>**(HCV genotype 3)** | N/A | 12 |

### Clinical Screening

Liver function tests and full blood counts for all 16 ESN women were within the normal ranges (Supplementary Table 3). The occurrence of autoimmune reactivity and autoimmune diseases within the ESN group was not more frequent than the seropositive group (Supplementary Table 3). However, many of the 16 ESN women had non-specific inflammatory symptoms including chronic fatigue (14/16; 88%) and muscular ache and pains consistent with a diagnosis of fibromyalgia (7/16; 44%), as detailed in Supplementary Table 3.

All ESN women were screened for anti-HCV antibodies using the ARCHITECT anti-HCV assay, and all tested negative. No statistically significant differences in HCV-specific T cells responses were observed between the ESN group and matched healthy controls (Figure 2A). HCV-specific T cell responses, greater than the mean+3SD of the HC group, were detected in two ESN individuals (Figure 2A). One ESN individual responded to the two NS3 peptide pools with 25 and 70 counts per 10^6^ PBMCs, respectively, and another ESN individual responded to the NS4A/B peptide pool with 58 counts per 10^6^ PBMCs. These responses were comparable in magnitude to HCV-specific T cell responses observed in seropositive spontaneous resolvers and infected individuals who achieved SVR following antiviral therapy (Figure 2A). HCV-specific T cell responses, in the absence of antibody reactivity, are a feature of published ESN cohorts(3–10). No neutralising antibody responses targeting the HCV E1/E2 glycoproteins were detected in the ESN cohort (Figure 2B). All ESN individuals were seropositive for anti-EBV while 4 (out of 16; 25%) were seropositive for anti-CMV. This low rate of CMV seropositivity has been previously observed in the Irish population although the reasons for this are unknown(25).

**Figure 2.**
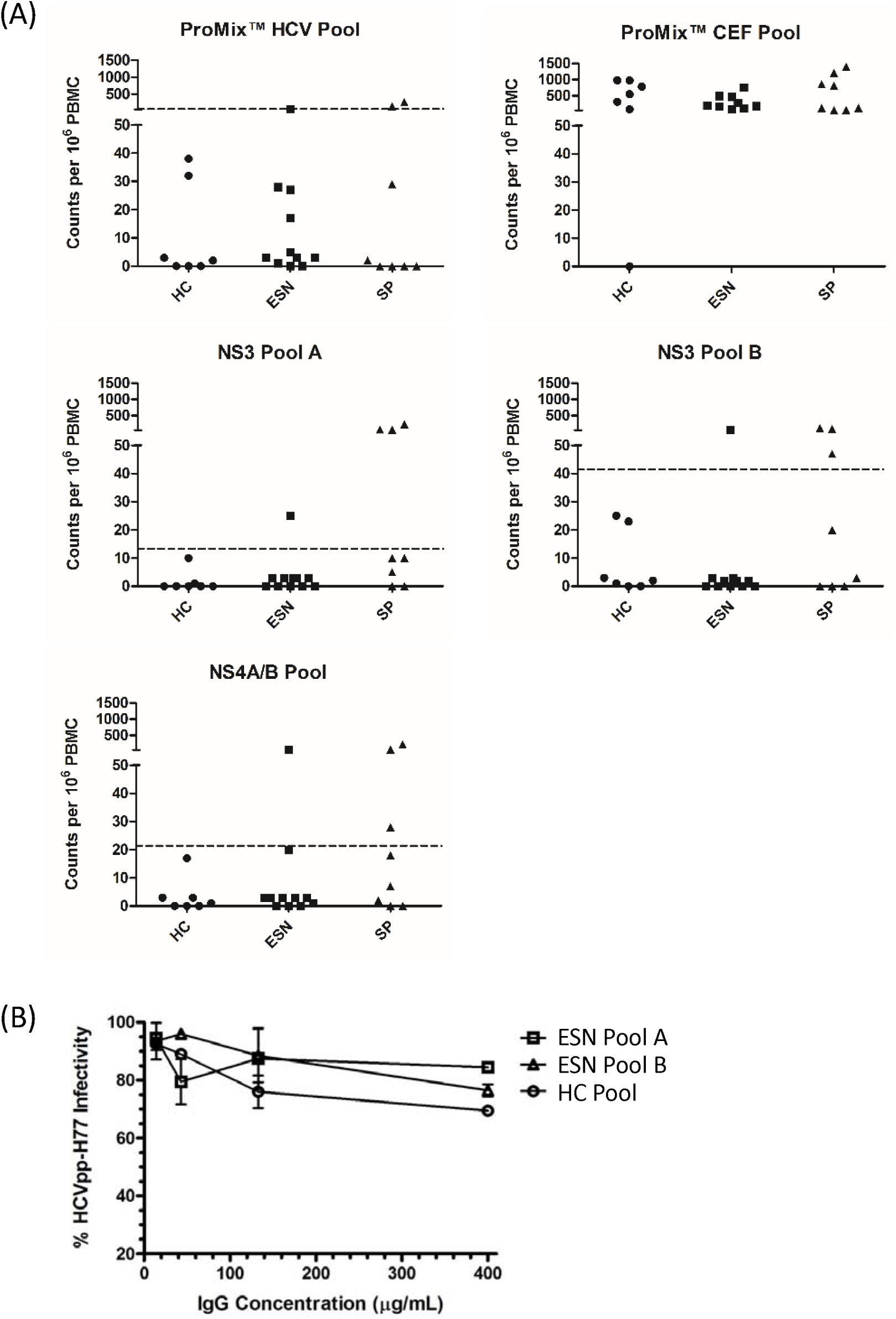
T cell IFNγ ELISpots and HCVpp-H77 Neutralisation. (A) IFNγ-producing T cell counts per 10^6^ PBMC in age- and gender-matched healthy controls (HC), seropositive (SP) individuals, and exposed seronegative (ESN) individuals following stimulation with ProMix™ HCV and CEF peptide pools and 13 to 19-mer peptide pools spanning NS3, NS4A, and NS4B (as detailed in Supplementary Table 1). The dashed line represents the mean + 3×SD of the HC group. (B) HCVpp-H77 infectivity, following incubation with purified IgG from pooled plasma from HC (n=9) and SN individuals (two pools each with n=8), to assess neutralising antibody responses targeting HCV E1/E2 glycoproteins. IgG samples were tested in triplicate and the results are the mean (and SD) of two independent experiments.

### Immune cell frequency and IFN*α* responsiveness

Amongst the ESN individuals a statistically significant decrease in the frequency of CD14+ monocytes was observed compared to age- and gender-matched controls (Figure 3A). No change in the frequency of T cells, B cells or NK cells, as a proportion of total leukocytes, was observed (Figure 3A). Type I IFNs are the key anti-viral effector molecules of the innate immune system. To assess immune cell responsiveness to type I IFNs, whole blood was stimulated with IFNα, and levels of STAT1 and STAT3 phosphorylation were detected after 30 minutes. No differences in STAT1 phosphorylation were observed (Figure 3B), however, ESN individuals showed enhanced STAT3 phosphorylation, specifically within their NK cells, compared to age- and gender-matched controls (Figure 3C). This was not due to a difference in basal STAT3 phosphorylation levels (Figure 3D) or STAT3 expression levels (data not shown).

**Figure 3.**
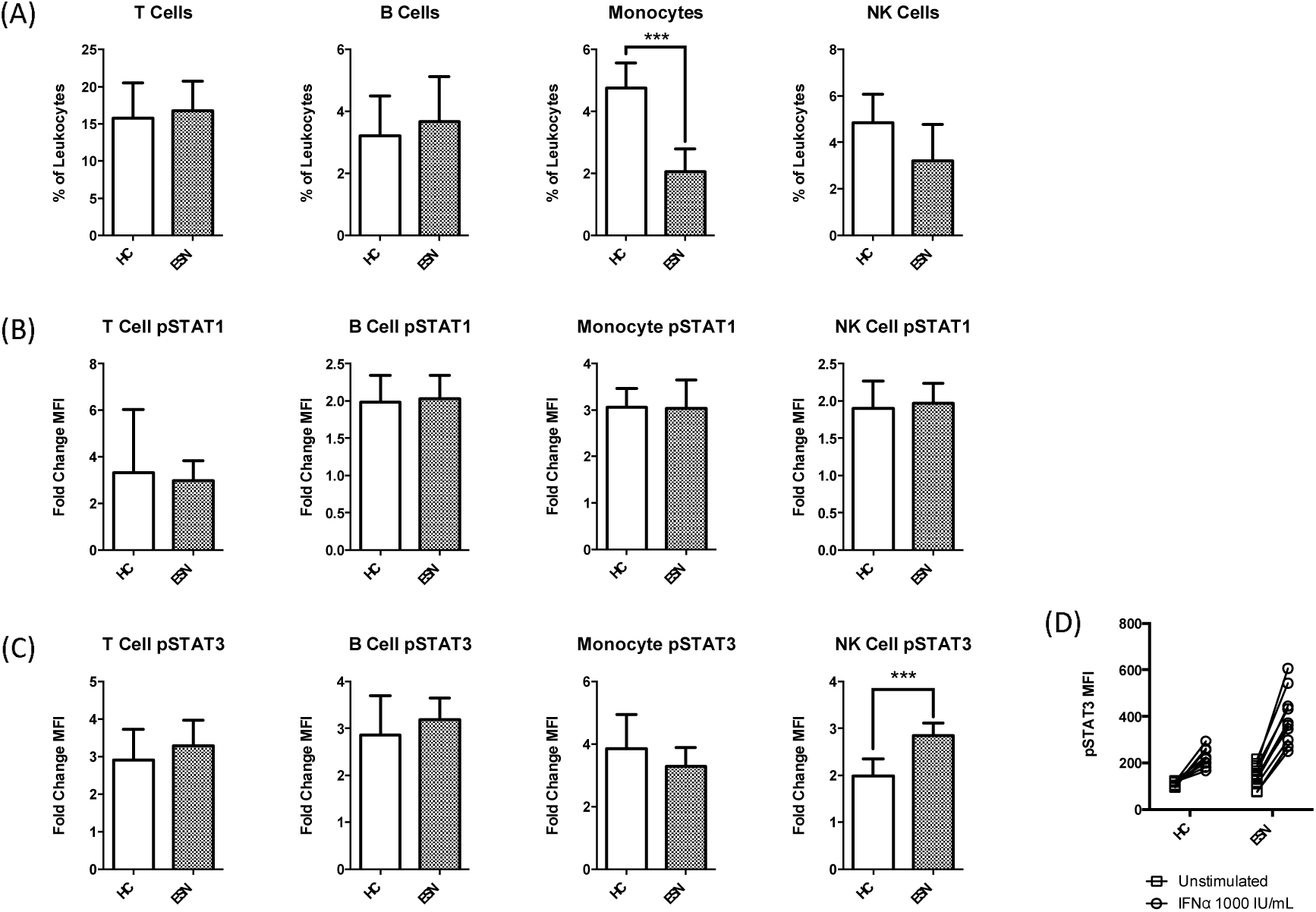
Leukocyte frequency and responsiveness to Type I IFN in age- and gender-matched health control (HC) and exposed seronegative (ESN) individuals. (A) Frequency of T cells, B cells, monocytes and NK cells as a percentage of total leukocytes. (B) Median fluorescence intensity (MFI) fold change of pSTAT1 in T cells, B cells, monocytes and NK cells, following 30 minutes stimulation with 1000 IU/mL IFNα. (C) Median fluorescence intensity (MFI) fold change of pSTAT3 in T cells, B cells, monocytes and NK cells, following 30 minutes stimulation with 1000 IU/mL IFNα. (D) MFI values for pSTAT3 in unstimulated and IFNα-stimulated NK cells. (A) Unpaired t-test and (B-C) two-tailed Mann-Whitney test; *** *P*<0.001.

### NK cell phenotype and function

To further explore the phenotype and function of NK cells within this cohort of ESN individuals, expression of cell surface receptors and functional assays were performed. No significant differences in the expression of the activation marker CD57 or NK receptors (NKG2A, NKG2D, NKp30, NKp44, and NKp46) were observed between age- and gender-matched healthy controls and ESN individuals (Supplementary Figure 2). To assess NK cell function in ESN individuals, PBMCs were stimulated overnight with cytokines and then co-incubated with 721.221 HLA-negative target cells. No differences were observed in CD107a expression (a marker of NK cell degranulation) between the age- and gender-matched healthy controls and the ESN individuals (Figure 4A and B). In contrast, IFNγ production was significantly increased within total NK cells from the ESN individuals following IL2/IL12/IL18 stimulation and target cell co-incubation (mean percentage IFNγ^+^ 50.1 vs 28.3 in healthy controls; Figure 4A). When NK cell subpopulations were analysed, increased IFNγ production was restricted to the CD56^dim^ NK cells (mean percentage IFNγ^+^ 48.3 vs 31.8 in healthy controls; Figure 4B).

**Figure 4.**
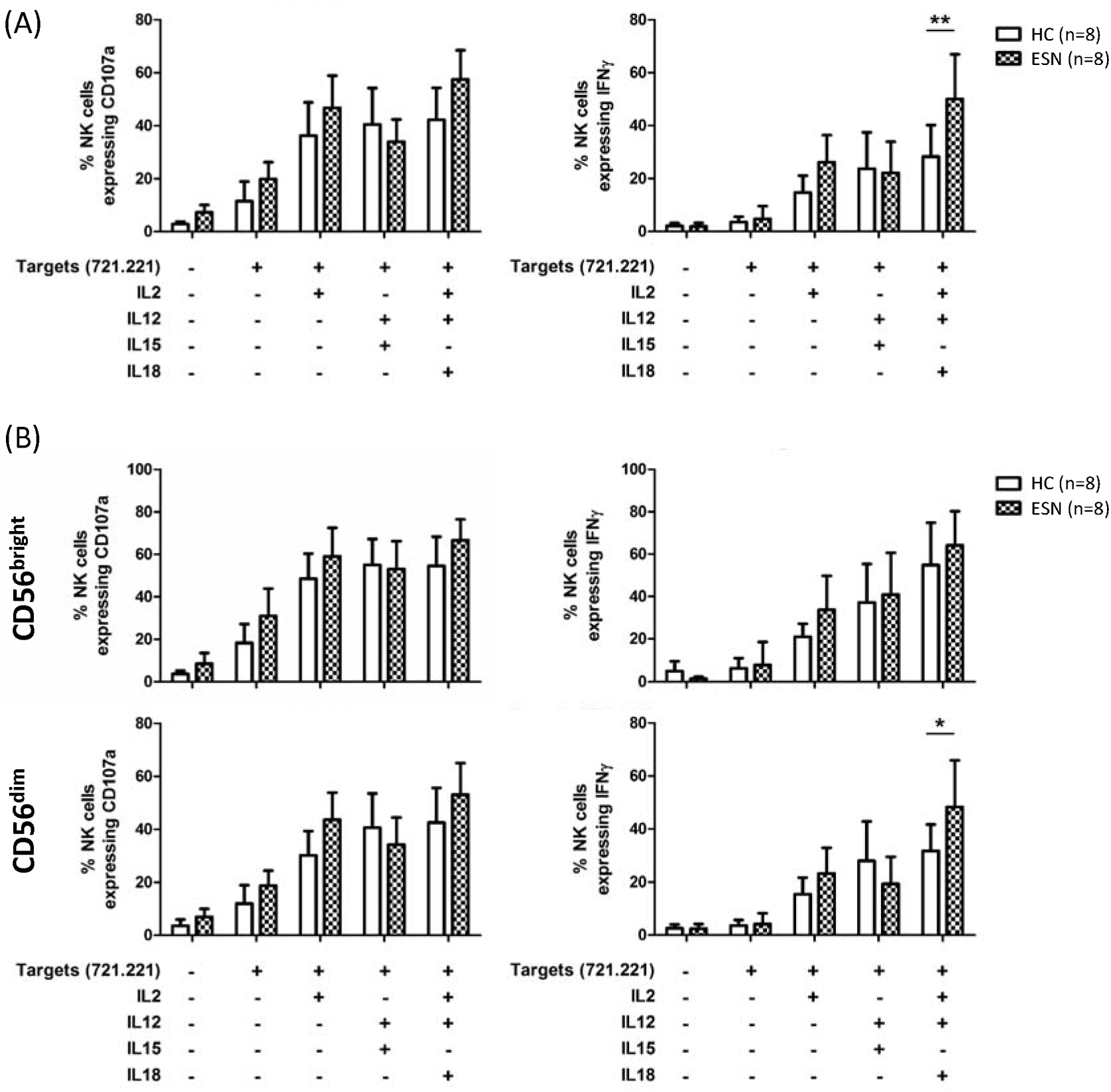
NK cell function in age- and gender-matched health control (HC) and exposed seronegative (ESN) individuals. (A) Percentage of total NK cells expressing IFNγ and the degranulation marker CD107a in HC (n=8) and SN (n=8) following cytokine and target cell stimulation. (B) Percentage of CD56^bright^ and CD56^dim^ NK cells expressing IFNγ and the degranulation marker CD107a in HC (n=8) and SN (n=8) following cytokine and target cell stimulation. Data presented as mean and 95% confidence interval. Two-way ANOVA with Bonferroni post-test; * *P*<0.05; ** *P*<0.01.

### Cirulating cytokine levels

To determine whether ESN individuals had elevated pro-inflammatory cytokine levels, plasma samples were analysed using the BioLegend LEGENDplex™ Human Inflammation Panel (13-plex). There was no evidence for a distinct pro-inflammatory cytokine signature in the ESN individuals, as demonstrated by the lack of clustering of ESN and healthy controls on the basis of cytokine levels (Figure 5A). Large inter-individual variation was observed amongst the ESN individuals, with two ESN individuals in particular having elevated levels of a range of pro-inflammatory cytokines (Figure 5A). ESN individuals did have significantly higher levels of circulating IL8 (mean 312.6 vs 47.1 ng mL^-1^ in healthy controls; Figure 5B), as well as higher levels of circulating IL18 (mean 405.4 vs 233.0 ng mL^-1^ in healthy controls; Figure 5B).

**Figure 5.**
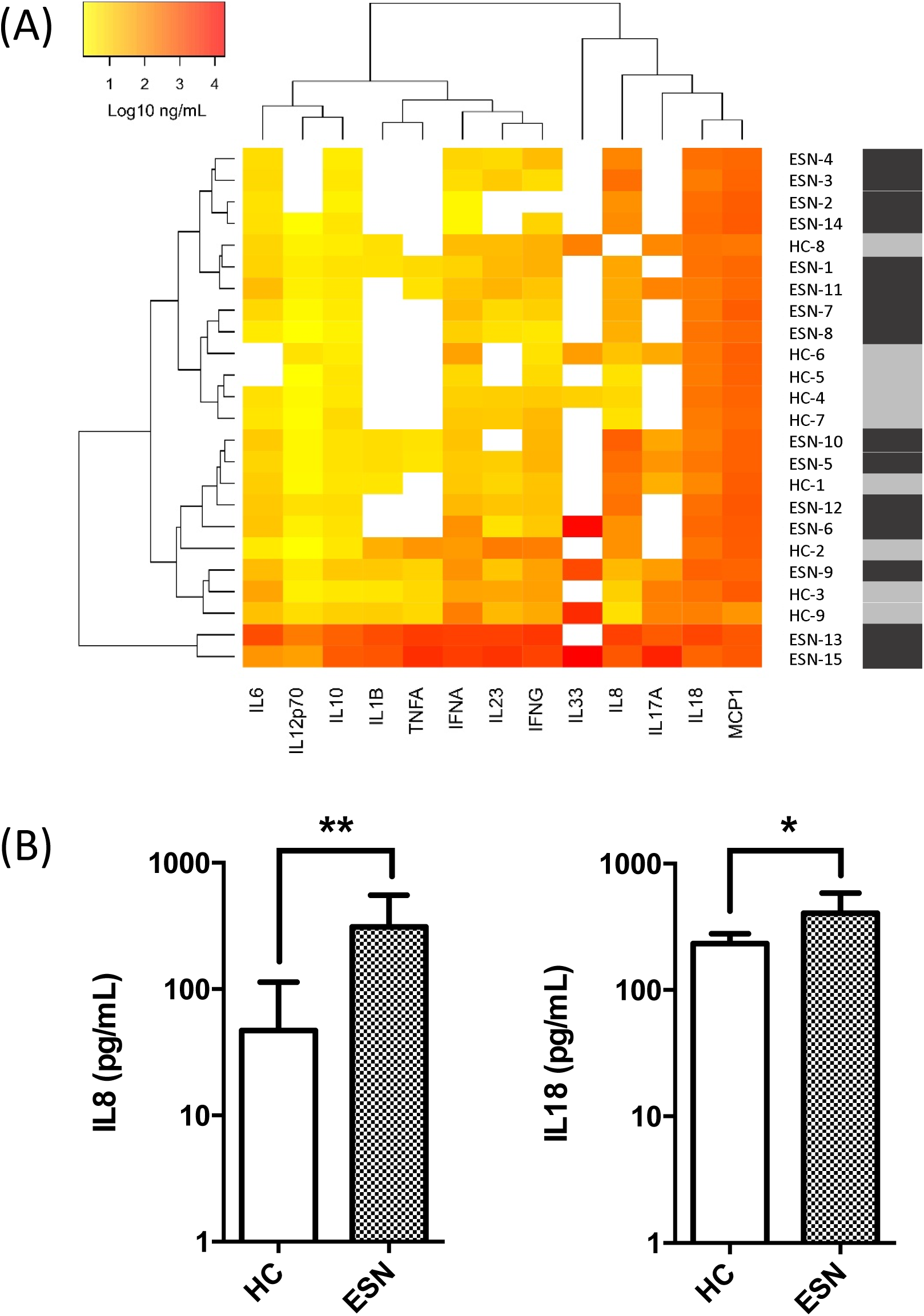
Plasma cytokine profiles in age- and gender-matched health control (HC) and exposed seronegative (ESN) individuals. (A) Heat map of plasma cytokine levels (log 10 ng mL^-1^) in HC (n=9) and SN (n=15). (B) Plasma IL8 and IL18 levels (ng mL^-1^) in HC (n=8) and SN (n=15). Data presented as mean and 95% confidence interval. Two-tailed Mann-Whitney test; * *P*<0.05; ** *P*<0.01.

## Discussion

People who are exposed to HCV infection but remain seronegative (ESN) provide opportunities to explore the intrinsic and innate immune pathways that confer resistance to HCV infection and possibly inform more generally about innate resistance to viral infection. ESN cohorts have been described for both HCV and HIV(26, 27). In both infections, ESN individuals possess enhanced innate immune function. Several of these cohorts are limited by variability in the viral strains between individuals, as well as uncertainty in the time of exposure. Here for the first time, we describe ESN individuals amongst Irish women who received potentially HCV-contaminated anti-D immunoglobulin between 1977-1979 or 1991-1994. These two single-source HCV outbreaks were associated with a single-source viral strain and involve a large population of women with defined dates of exposure(20, 21).

The ESN women in the present study had no antibody-responses specific for HCV at the time of the present study. HCV screening for anti-D recipients was first initiated in Ireland in 1994, meaning individuals were first screened 17 years following the start of the first anti-D contamination period, and 3 years following the start of the second anti-D contamination period. It is possible that some ESN individuals had developed HCV-specific antibodies but were seronegative at the date of screening due to the natural attrition rate of specific antibodies in individuals who spontaneously resolve HCV infection(28). HCV-specific T cell responses have also been shown to wane within months of cessation of viral exposure in ESN individuals(9), although they are thought to persist longer than HCV-specific antibody responses(28). In the present study, two individuals possessed T cell responses specific for HCV non-structural proteins, which were greater than 3SD above the mean of an age- and gender-matched healthy control group.

The NK cells in our ESN group showed enhanced IFNγ production following cytokine stimulation. Enhanced IFNγ production in total NK cells as well as CD56^dim^ NK cells has been previously observed in ESN individuals following exposure(13, 15). These results suggest that enhanced NK cell IFNγ production may contribute to the early clearance of HCV, although this presumes that the enhanced IFNγ NK cell phenotype is stable over time, which is currently unknown. Our results support the idea that a spectrum of NK cell function exists within the unexposed healthy population, and following exposure individuals with high NK cell function are more likely to rapidly clear the virus.

In addition to enhanced NK cell responses, we observed significantly elevated circulating levels of the chemokine IL8, as well as increased IL18. IL18 directly primes NK cells and promotes IFNγ production upon NK cell activation(29), possibly providing a link to the enhancement of NK cell functions. IL8 is a chemokine involved in neutrophil recruitment and a previous study also identified elevated circulating IL8 levels in ESN individuals(30). While neutrophils are commonly associated with bacterial clearance, they also play important antiviral roles(31) and the release of neutrophil extracellular traps can protect against viral challenge(32).

The classification of individuals as ESN in our study is complicated by a number of factors. A lack of HCV-specific antibodies upon HCV exposure may be due to different biological pathways, which confer intrinsic (entry receptor mutation), innate (NK cell-mediated), or adaptive (T cell-mediated) immune resistance to HCV infection, and these pathways may be different between ESN individuals. Secondly, it is possible that some individuals appear to be ESN due to exposure to negligible doses of HCV. Inter- and intra-batch variation in infectious viral load is an unavoidable confounding factor in assigning exposure risks within ESN cohorts. In the absence of detectable HCV-specific adaptive immune responses it is impossible to confirm or exclude past viral exposure. The self-reported nature of our ESN cohort may also influence our data, in particular the high prevalence of non-specific apparent inflammatory symptoms, including chronic fatigue and muscular ache and pains consistent with a diagnosis of fibromyalgia. The heterogeneity within our ESN classification is a limitation of this pilot study.

A better understanding of the genetic and immune parameters that confer natural resistance to viral infection is required for the development of novel strategies to combat emerging and re-emerging viral outbreaks. Well-defined human ESN cohorts provide unique insights into the biological mechanisms associated with natural resistance to viral infection and can identify these antigen-independent processes that effectively block viral infection within resistant members of the human population.

## Acknowledgements and Disclosures

We wish to thank the volunteers who took part in this study, the patient support group Positive Action, and in particular Evelyn Quinn for her help. We also wish to thank Dr Louise Pomeroy and Carmel Sheridan for their assistance. The peptides detailed in Supplementary Table 1 were kindly obtained through BEI Resources, NIAID, NIH. This study was funded by a HRB grant to CG (HRB.POR/2011.16) and a Science Foundation Ireland Investigator Award to COF (12/IA/1667).

