## Supplementary for "Enhanced cytokine responsiveness in natural killer cells from a pilot cohort of uninfected seronegative women exposed to hepatitis C virus contaminated anti-D immunoglobulin"

|  | **HCV Genotype** | **GenPept Accession** | **BEI Resources Catalog No.** | **HCV Protein** | **Peptide Numbers** | **Amino Acid Coverage** |
| --- | --- | --- | --- | --- | --- | --- |
| **NS3 Pool A; J4** | 1b | AAC15722 | NR-3742 | NS3 | 1-45 | 1-293 |
| **NS3 Pool B; J4** | 1b | AAC15722 | NR-3742 | NS3 | 46-98 | 283-631 |
| **NS4A/B Pool; J4** | 1b | AAC15722 | NR-3743  NR-3744 | NS4A  NS4B | 1-7  1-40 | 1-54  1-261 |
| **NS3 Pool A; K3a/650** | 3a | BAA06044 | NR-4066 | NS3 | 1-45 | 1-300 |
| **NS3 Pool B; K3a/650** | 3a | BAA06044 | NR-4066 | NS3 | 46-97 | 290-631 |
| **NS4A/B Pool; K3a/650** | 3a | BAA06044 | NR-4067  NR-4068 | NS4A  NS4B | 1-7  1-39 | 1-54  1-261 |

**Supplementary Table 1: Peptide pools used for ELISpot assays.**

| Target | Fluorophore | Clone | Supplier | Cat. No. |
| --- | --- | --- | --- | --- |
| CD107a | APC | H4A3 | BioLegend | 328619 |
| CD11c | eFluor-450 | 3.9 | eBioscience | 48-0116-42 |
| CD123 | PE-Cy7 | 6H6 | eBioscience | 25-1239-42 |
| CD14 | V450 | M5E2 | BD Biosciences | 561390 |
| CD16 | PE-Cy7 | 3G8 | BD Biosciences | 560918 |
| CD16 | FITC | B73.1 | BD Biosciences | 561308 |
| CD19 | APC-H7 | HIB19 | BD Biosciences | 560727 |
| CD3 | V500 | UCHT1 | BD Biosciences | 561416 |
| CD3 | BV421 | UCHT1 | BD Biosciences | 562427 |
| CD3 | BV605 | OKT3 | BioLegend | 317321 |
| CD45 | BV510 | HI30 | BD Biosciences | 563204 |
| CD56 | PE-Cy7 | HCD56 | BioLegend | 318318 |
| CD56 | A488 | B159 | BD Biosciences | 557699 |
| CD56 | BV711 | NCAM16.2 | BD Biosciences | 563169 |
| CD57 | FITC | HCD57 | BioLegend | 322306 |
| HLA-DR | V500 | G46-6 | BD Biosciences | 561224 |
| IFNγ | PerCP-Cy5.5 | 4S.B3 | BioLegend | 502525 |
| Lin1 | FITC | SK7, 3G8, SJ25C1, L27, MφP9, NCAM16.2 | BD Biosciences | 340546 |
| NKG2A | PE-Cy7 | Z199 | Beckman Coulter | A60797 |
| NKG2D | PE-CF594 | 1D11 | BD Biosciences | 562498 |
| NKp30 | APC | AF29-4D12 | Miltenyi Biotec | 130-099-380 |
| NKp44 | AF488 | 3.43 | AbD Serotech | MCA4665 |
| NKp46 | BV786 | 9E2 | BD Biosciences | 563329 |
| pSTAT1 (pY701) | A647 | 4a | BD Biosciences | 612597 |
| pSTAT3 (pY705) | PE | 4/P-STAT3 | BD Biosciences | 612569 |

**Supplementary Table 2: Flow Cytometry Antibodies.**

|  | **Exposed Seronegative** | **Seropositive** |
| --- | --- | --- |
| **n** | 16 | 10 |
| **Age (Years)** | 57.8 (52.4 - 61.3) | 62.5 (58.0 - 67.1) |
| **No. Exposed 1977-79 (HCV genotype 1A)** | 5 | 8 |
| **No. Exposed 1991-94 (HCV genotype 3)** | 12 | 2 |
| **No. Spontaneous Resolver** | N/A | 5 |
| **No. Treatment SVR** | N/A | 5 |
| **Body Mass Index** | 26.2 (23.4-29.0) | 23.9 (19.7-28.1) |
| ***Clinical History*** |  |  |
| **Chronic Fatigue** | 14 (16)** | 3 (10) |
| **Muscular Ache and Pains** | 7 (16) | 1 (10) |
| **Diabetes** | 1 (16) | 2 (10) |
| **Thyroid Disease** | 2 (16) | 2 (10) |
| **Kidney Disease** | 0 (16) | 0 (10) |
| **Liver Disease** | 0 (16) | 0 (10) |
| **Hypertension** | 5 (16) | 2 (10) |
| **Osteoarthritis** | 6 (16) | 1 (10) |
| **Asthma** | 2 (16) | 1 (10) |
| **Heart Disease** | 0 (15) | 1 (10) |
| **Cancer** | 1 (16) | 0 (10) |
| **Current Smokers** | 2 (16) | 3 (10) |
| **Ever Smokers** | 5 (16) | 6 (10) |
| ***Immunology Tests*** |  |  |
| **Anti-Nuclear Antibody** | 4 (14) | 5 (10) |
| **Anti-Thyroid Peroxidase Antibody** | 0.3 (0.0-47.5) | 0.7 (0.1-415.3) |
| **Anti-Tissue Transglutaminase Antibody** | 0.8 (0.3-4.7) | 0.4 (0.1-1.9) |
| **Anti-Smooth Muscle Antibody** | 0 (16) | 0 (9) |
| **Anti-Parietal Cell Antibody** | 3 (16) | 1 (9) |
| **Anti-Mitochondrial Antibody** | 0 (16) | 0 (9) |
| **Anti-Liver Kidney Microsomal Antibody** | 0 (16) | 0 (9) |
| **IgM Rheumatoid Factor** | 0.5 (0.4-22) | 0.7 (0.4-6.5) |
| ***Biochemical Tests*** |  |  |
| **Albumin (g/L)** | 40.1 (38.8-41.4) | 39.1 (37.5-40.7) |
| **Total Bilirubin (µmol/L)** | 7.6 (6.1-9.1) | 5.6 (4.0-7.2) |
| **Alkaline Phosphatase (U/L)** | 77.6 (67.5-87.8) | 79.0 (71.1-86.9) |
| **Gamma-Glutamyl Transferase (U/L)** | 31.4 (18.1-44.6) | 23.6 (15.9-31.3) |
| **Alanine Aminotransferase (U/L)** | 25.6 (19.0-32.1) | 23.6 (13.7-33.5) |
| **Aspartate Aminotransferase (U/L)** | 23.7 (20.0-27.4) | 23.0 (16.4-29.6) |
| **White Cell Count (10^9^/L)** | 6.6 (5.8-7.3) | 7.8 (6.4-9.3) |

**Supplementary Table 3. Cohort demographics and clinical data.** Data for age, body mass index and biochemical & haematology tests are presented as mean (95% confidence intervals). Data for clinical history are presented as total number of cases (total number assessed). Data for immunology tests are presented as number of positives (total number assessed) or as median (range). ** *P*-value <0.01, Chi-Square Test; *** *P*-value <0.001, unpaired t-test.

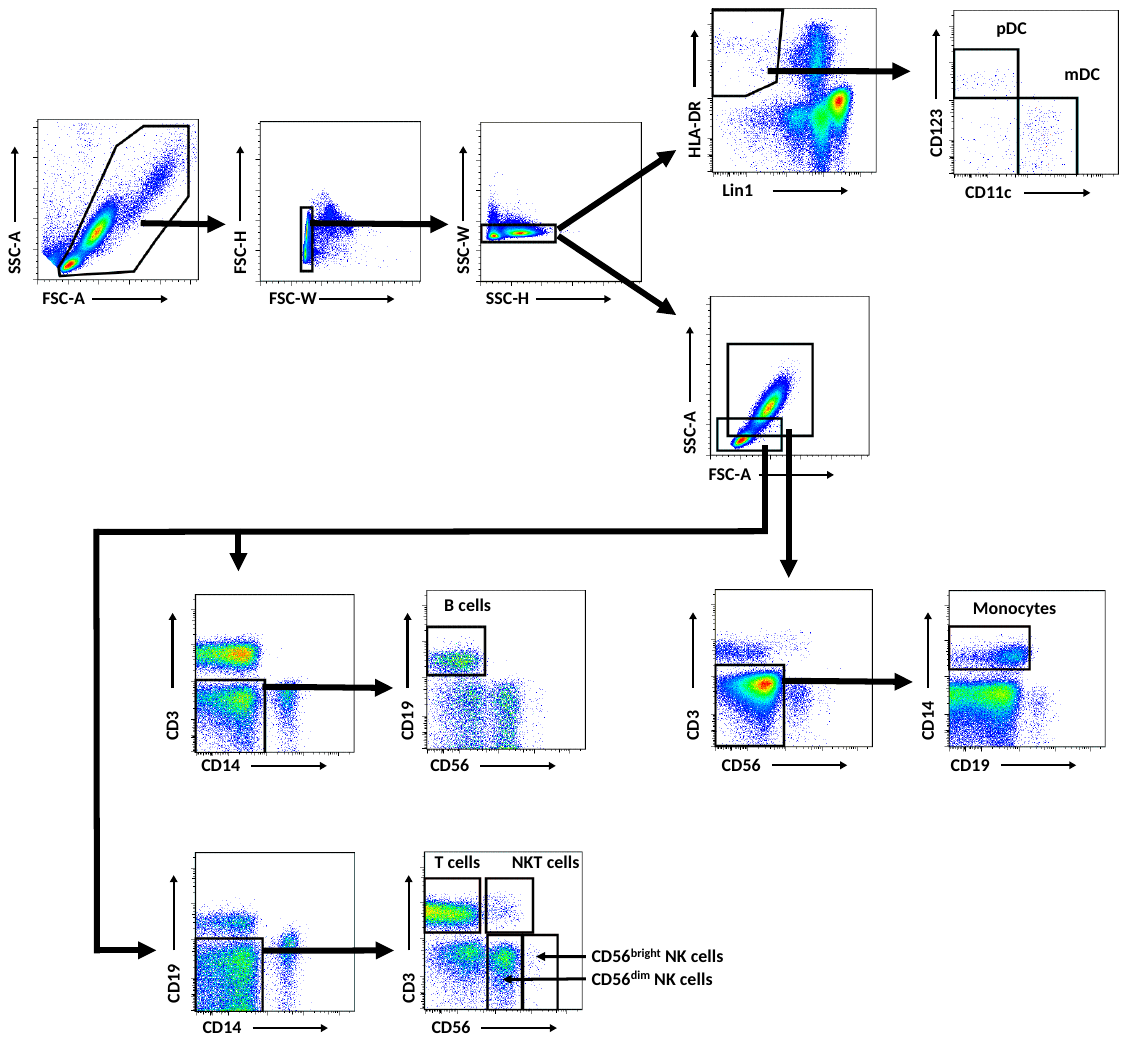

**Supplementary Figure 1. Representative gating strategy for flow cytometry analysis.**

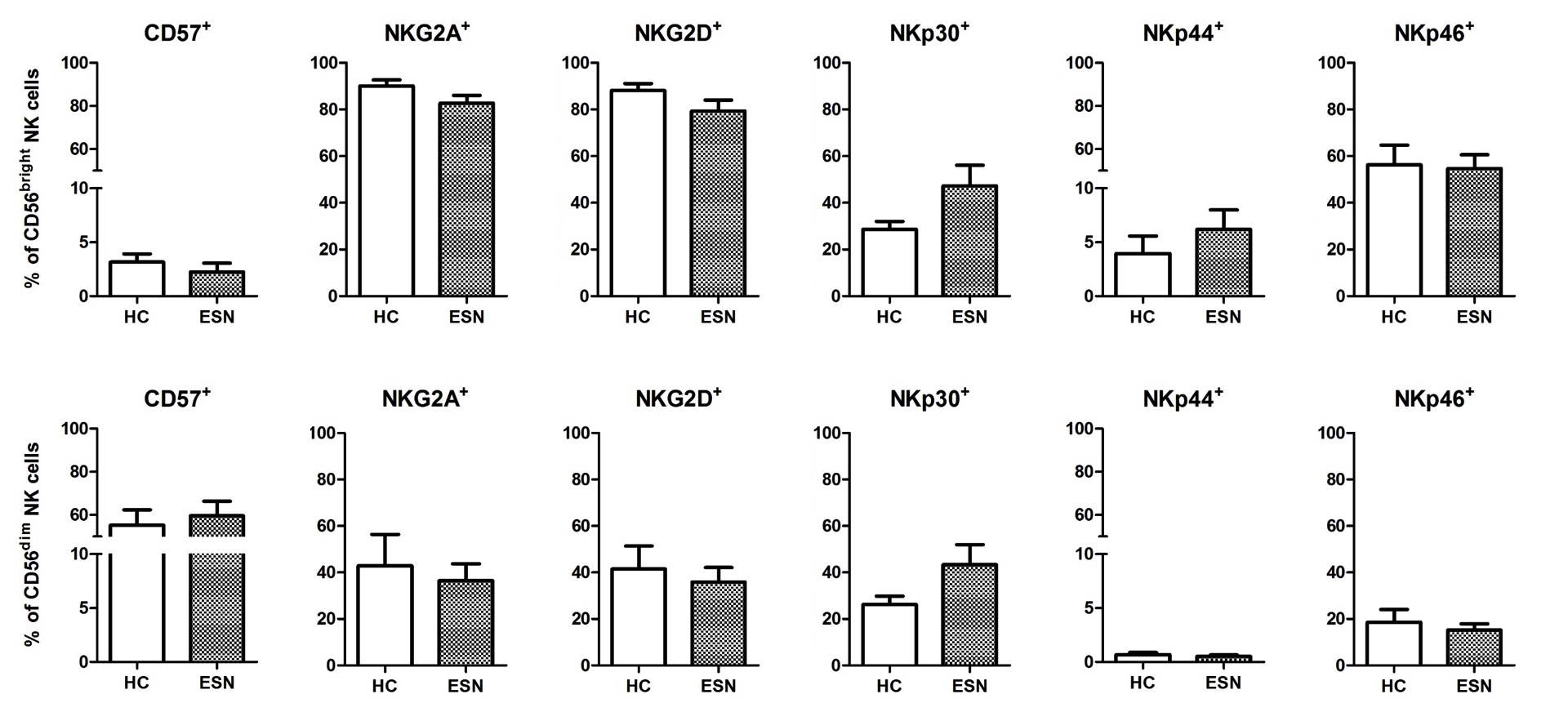

**Supplementary Figure 2. NK cell phenotype in age- and gender-matched health control (HC) and exposed seronegative (ESN) individuals.** Percentage of CD56^bright^ CD16^-^ NK cells (top panels) and CD56^dim^ CD16^+^ NK cells (bottom panels) expressing cell surface markers comparing HC (n=4, except for CD57 where n=8) and ESN (n=7, except for CD57 where n=8). Data presented as mean and 95% confidence interval.
